# Gene networks underlying faster flowering induction in response to far-red light

**DOI:** 10.1101/234161

**Authors:** Maria Pazos-Navarro, Federico M Ribalta, Bhavna Hurgobin, Janine S Croser, Parwinder Kaur

## Abstract

Light is one of the main signals that regulates flowering. Low red to far-red ratios accelerate flowering in a wide range of species. The central gene pathways controlling flowering time in Arabidopsis, appear to be largely conserved in legumes. However, numerous examples exist of gene duplication and loss. The role of *CONSTANS-LIKE* genes as integrators of the photoperiod response has been questioned in several dicot species, including legumes. In this study on subterranean clover, using RNA-seq and controlled light spectra, we identified 13 differentially expressed genes related to light signalling, meristem identity and flowering promotion. Of these, we pinpointed genes which seem to link photoperiod and far-red light signalling coding for a With no lysine kinase, a CCT motif related to *CONSTANS*, a *FLOWERING LOCUS T b2* like protein, and their active downstream cascade. The earlier down-regulation of these genes under blue compared to far-red-enriched light may explain their role in floral induction. A second independent approach (qPCR analysis) confirmed our findings. Contrasting responses to light quality related to reproduction and defence mechanisms were also found. These results will contribute to a better understanding of the molecular basis of flowering in response to light quality in long-day plants.

## INTRODUCTION

Flowering behaviour is modified by environmental cues, with light being one of the most important signals that regulates flowering through quality, quantity and duration (Thomas 2006). Plants perceive subtle changes in light composition, duration and direction through photoreceptor pigments, which result in physiological and morphological modifications necessary for adaptation to environmental changes (Rajapakse & Shakak 2007). In higher plants, the ‘shade avoidance syndrome’ is an adaptive developmental strategy mediated by the perception of light spectral quality, in particular the red to far-red ratio, R: FR, which acts as an early warning of potential shading (Salter *et al.* 2003; Ballaré & Pierik 2017). In Arabidopsis (*Arabidopsis thaliana*), the most noticeable shade avoidance responses include rapid internode elongation and accelerated floral initiation at the expense of leaf expansion and chlorophyll synthesis reduction (Devlin *et al.* 1999). Similarly, research in the model long-day species pea (*Pisum sativum* L.) demonstrates FR-enriched light, with a R: FR ratio below 3.5, is most effective for early floral induction (Runkle & Heins 2001; Cummings *et al.* 2007; Croser *et al.* 2016; Ribalta *et al.* 2017).

Changes in light quality are detected in the leaf by the action of a family of plant photoreceptors, including phytochromes (R and FR light receptors) and cryptochromes (blue light receptors) and involve complex gene regulatory networks (Andrés & Coupland 2012; Viczián *et al.* 2017). In Arabidopsis significant progress has been made toward understanding the role light quality plays on floral initiation pathways. The genes *GIGANTEA* (*GI*), *FLAVON KELCH F BOX* (*FKF1*), *CONSTANS* (*CO*) and *FLOWERING LOCUS T* (*FT*) have major regulatory roles in the promotion of flowering in response to photoperiod (Thomas 2006; Jiao *et al.* 2007). In particular, *CO* has been described as a network hub for the integration of internal and external signals into the photoperiodic flowering pathway to induce *FT* expression (Wong *et al.* 2014; Shim *et al.* 2017). *FT* acts as a mobile floral promoting signal that integrates day length, light quality, circadian clock, temperature and vernalisation inputs (Turck *et al.* 2008). In Arabidopsis, the role of the circadian clock and light on the regulation of *CO* gene expression is well understood (Imaizumi & Kay 2006; Cerdan & Chory 2003). The circadian clock sets *CO* expression in the afternoon in long-day plants and light activates CO protein to induce *FT* (Imaizumi & Kay 2006).

In legumes, the genes and gene families central to pathways controlling flowering time in Arabidopsis appear to be largely conserved. However, there are numerous examples of gene duplication and loss (Weller & Ortega 2015). In pea, the conserved role of the Arabidopsis genes *GI, EARLY FLOWERING 3* and *EARLY FLOWERING 4* in the regulation of *FT* genes has been demonstrated (Hecht *et al.* 2007; Liew *et al.* 2009; Weller *et al.* 2012). Orthologous Arabidopsis genes play a part in the photoperiodic flowering pathway in legumes. However, the role of *CONSTANS-LIKE* (*COL*) genes as integrators of the photoperiod response has been questioned in several dicot species including legumes (Blackman 2017). Recent studies in *Medicago truncatula* Gaertn. (Medicago) revealed that none of the *COL* genes identified are functionally equivalent to *CO*, with respect to inducing *FT* expression (Wong *et al.* 2014). These findings support the idea of *CO*-independent pathways involved in flowering induction in legumes.

An improved understanding of the gene networks underlying flowering induction in response to light quality will require better characterisation of the transcriptome. The recent development of deep-sequencing technologies, such as RNA-Seq, has enabled the generation of a high-resolution global view of the transcriptome and its organisation for some species and cell types (Wang *et al.* 2009). Whole-transcriptome sequencing using RNA-Seq is a convenient and rapid means to study gene expression at the whole-genome level and define putative gene function (Wang *et al.* 2009; Ozsolak & Milos 2011; Singh *et al.* 2013; Hirakawa *et al.* 2016; Kaur *et al.* 2017). Rapid advances have been made toward understanding the transcriptional regulation of specific developmental processes in legumes (Benedito *et al.* 2008; Libault *et al.* 2008; Severin *et al.* 2010). Within this framework, we now seek to apply whole-transcriptome sequencing to characterise the genetic regulatory mechanisms underlying the induction of flowering in legumes in response to changes in the R: FR ratio using light emitting diodes (LED).

Despite the substantial number of grain and pasture legumes with their genome sequenced and/ or significant genomic resources such as chickpea (*Cicer arietinum* L.), pea, soybean (*Glycine max* (L). Merr.), Medicago and *Lotus japonicus*, no legume species has yet emerged as a predominant model in the study of flowering time (Weller & Ortega 2015). Recently, we established subterranean clover (*Trifolium subterraneum* L.) as a reference species for genetic and genomic studies within the genus *Trifolium*. Subterranean clover is diploid (2n = 16), predominantly inbreeding, and has a well-assembled and annotated genome with a tissue type transcriptome atlas available (Hirakawa *et al.* 2016; Kaur *et al.* 2017). The improved understanding of the genetics of flowering in this model species can assist to identify floral promotion pathways in other genetically complex species.

Within this research, we used deep-sequencing technologies to investigate transcriptional activity under different light spectra at three time-points in the long-day plant, subterranean clover. We have demonstrated accelerated onset of flowering across a range of leguminous species in response to FR-enriched LED light spectra (Croser *et al.* 2016; Pazos-Navarro *et al.* 2017; Ribalta *et al.* 2017). To elucidate the role of FR enriched spectra on acceleration of time to flowering, we have compared plants grown under a FR enriched regime with plants grown under a blue-enriched LED spectra. We expect that the FR-enriched spectra will accelerate the up-regulation of genes related to floral initiation.

## MATERIALS AND METHODS

### Experimental design

Seeds of *Trifolium subterraneum* L. cv. Daliak were nicked with a scalpel prior to sowing in 70 mm plastic pots filled with steam-pasteurised potting mix (Plant Bio Mix – Richgro Garden Products Australia Pty Ltd). Plants were grown simultaneously within two walk-in phytotron rooms under tightly controlled temperatures of 24 °C day, 20 °C night and a photoperiod of 20 h, as per Ribalta *et al.* (2017). The two growth environments differed only in the spectral composition of illumination provided to the plants (Fig. 1, A and B). Environment 1 (E1) illumination was provided by LED arrays enriched in the far-red part of the spectrum with a red to far-red (R: FR) ratio of 2.9 (‘B series’ AP67 spectrum Valoya Helsinki, Finland). Environment 2 (E2) illumination was provided by LED tubes enriched in the blue part of the spectrum with an R: FR ratio of 5.9 (108D18-V12 tubes from S-Tech Lighting, Australia). Spectral measurements were undertaken using a Sekonic C7000 SpectroMaster spectrometer (Sekonic Corp., Tokyo, Japan) and values were averaged over three scans in the range of 380–780 nm. Ratio calculations followed the method of Runkle & Heins (2001) where the R: FR ratio was measured as a narrow band (655–665: 725–735 nm). Plants were watered daily and fertilised weekly with a water-soluble NPK fertiliser (15:2.2:12.4) with micronutrients (Peters Excel, Scotts Australia, Bella Vista, New South Wales) at a rate of 2 g per pot, as per Pazos-Navarro *et al*. (2017). Time to flowering was defined as the number of days from sowing to the open floral stage of the first flower and was recorded under the different growth conditions.

**Figure 1.**
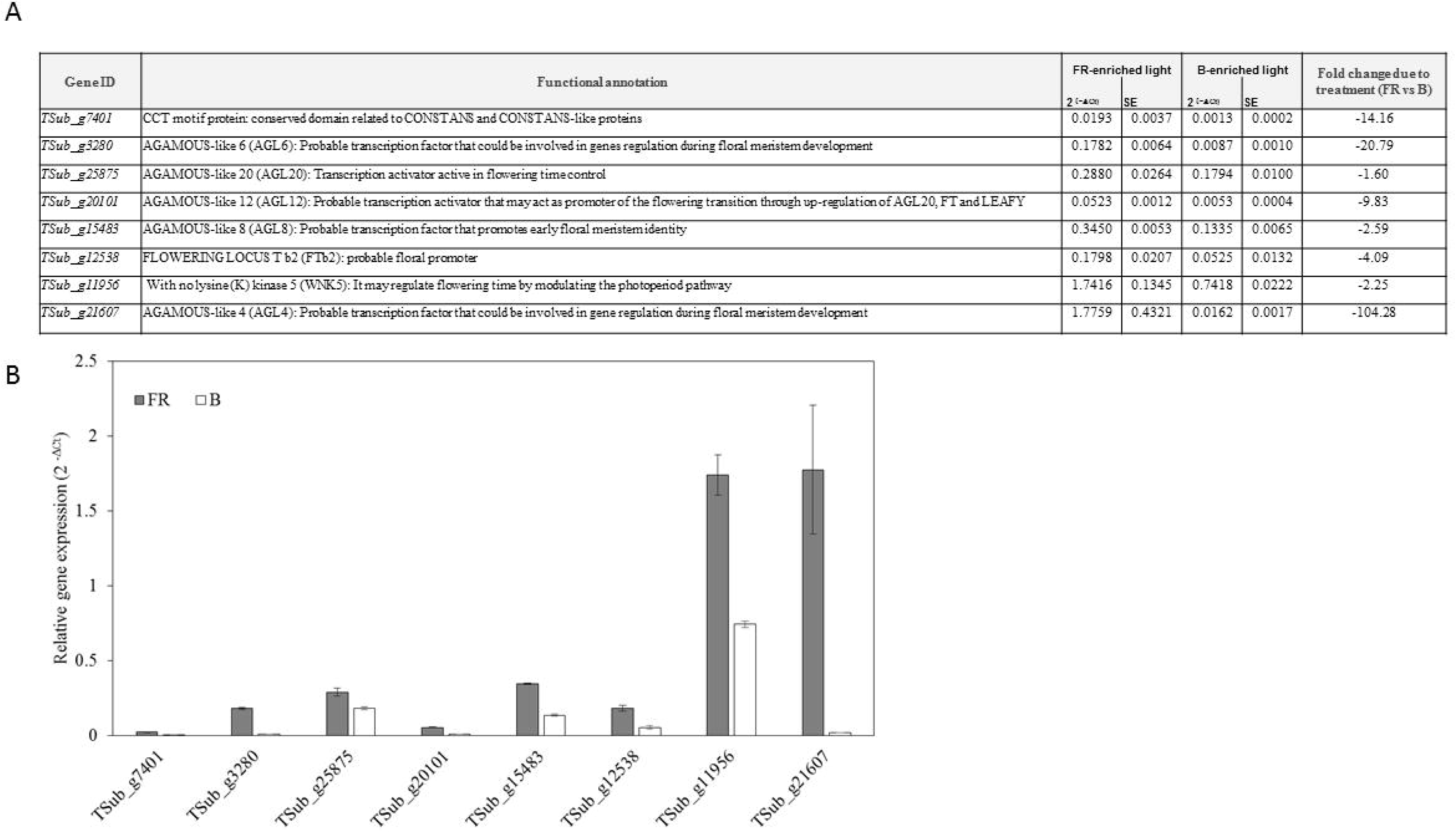
A. Light spectrum profiles (wavelength (nm) vs. photon flux density (PFD, µmol m^−2^ s^−1^). B. Time to flowering of *Trifolium subterraneum* plants and spectral characteristics (PFD, µmol m^−2^ s^−1^) of the environments used in this study. E1: far-red enriched light (AP67 LED lights, Valoya, Finland); E2: blue-enriched LED light (108D18-V12 tubes, S-Tech Lighting, Australia). The red to far-red ratio calculations followed the method of Runckle and Heins (2001): photon irradiance between 655 and 665 nm/ photon irradiance between 725and 735 nm.

Leaf samples were collected simultaneously 6 h after lights were switched on from E1 and E2 grown plants at three time-points (TPs) when plants in the E1 environment reached the following developmental milestones: third-leaf stage (TP1), appearance of the first flower bud (TP2) and open flower (TP3; Fig. 2A). This correlated to 14 growing days for TP1 (at this time-point, plants in both environments were at the third-leaf stage), 42 growing days for TP2, and 47 growing days for TP3. Leaf tissue (75–100 mg FW) was harvested and snap-frozen in liquid nitrogen and stored at –80 °C until RNA extraction.

**Figure 2.**
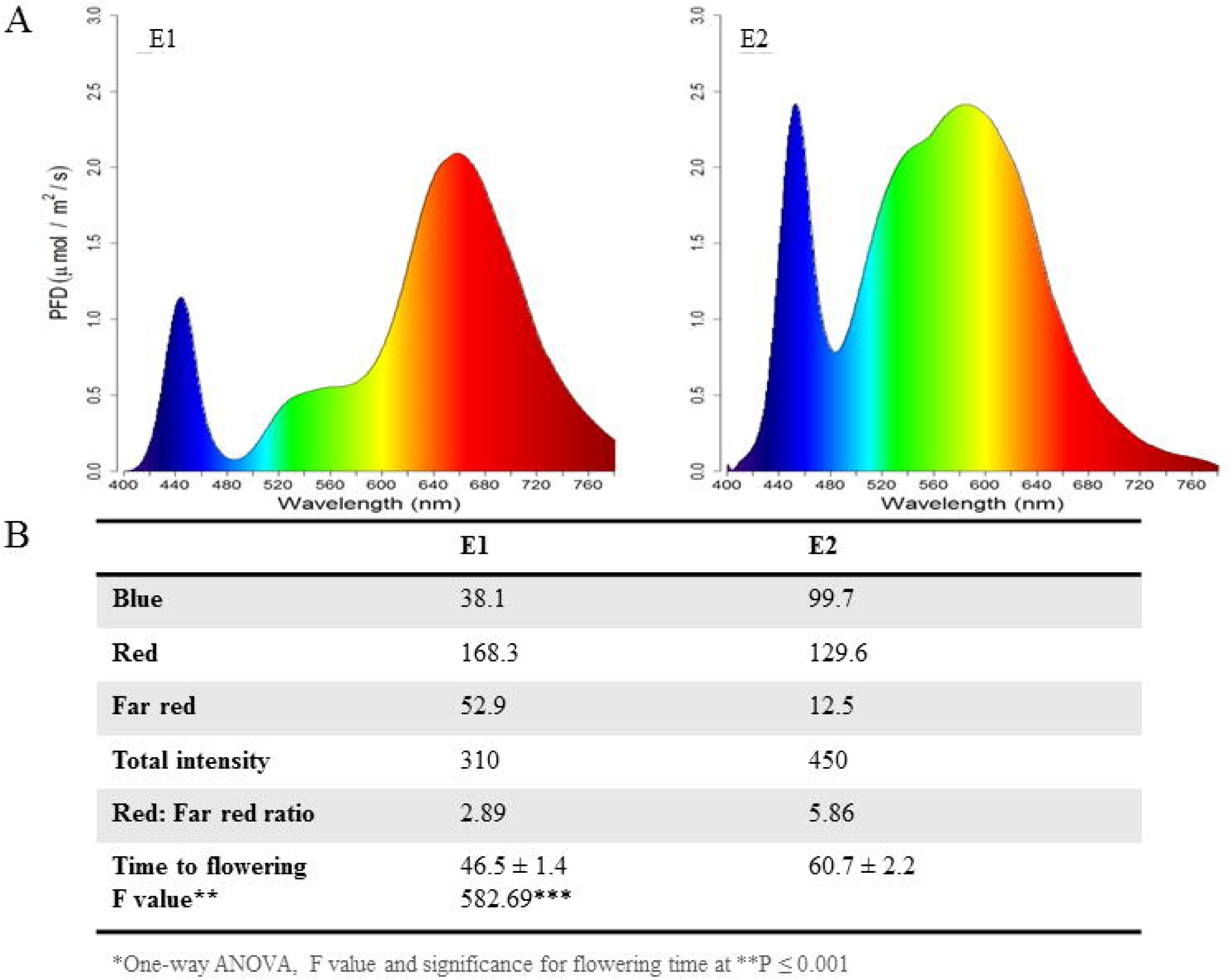
A. Time-points (TP) selected for this study based on when plants in the E1 environment attained the following developmental milestones: third-leaf stage (TP1), appearance of the first flower bud (TP2), open flower (TP3). B. Volcano plots for the differentially expressed genes. The red dots represent genes with logFC > 2 [FDR < 0.05] whereas the green dots represent genes logFC > 4 [FDR < 0.05].

### RNA isolation and library preparation

Total RNA from all tissue samples was extracted using the Spectrum(tm) Plant Total RNA Kit (Sigma-Aldrich, St Louis, USA) following the manufacturer’s instructions. Aliquots of purified RNA were stored at −80 °C. The concentration of RNA was confirmed using a Qubit fluorometer with the Qubit RNA assay kit (Life Technologies, Carlsbad, USA). The integrity of total RNA was determined by electrophoretic separation on 1.2% (w/v) denaturing agarose gels. Sequencing libraries were constructed using 500 ng of total RNA with a TruSeq^®^ Stranded Total RNA Sample Prep Kit with Ribo-Zero (Illumina Inc., San Diego, USA) following the manufacturer’s instructions. Concentrations of libraries were measured using the Qubit fluorometer with the Qubit dsDNA HS assay kit (Life Technologies, Carlsbad, USA) and Agilent high-sensitivity DNA chips (Agilent Technologies, Santa Clara, USA). The amplified libraries were pooled in equimolar amounts, and quality was assessed with Agilent high-sensitivity DNA chips (Agilent Technologies, Santa Clara, USA). All reads were pair-end sequenced using the HiSeq 2000 platform (Illumina Inc., San Diego, USA).

### Differential gene expression analysis

The sequencing quality of the Illumina reads was assessed using FastQC (Andrews 2010). The reads were quality filtered using Trimmomatic (Bolger *et al.* 2014) to remove adapter sequences. Filtered reads were mapped to the advanced genome assembly (Tsub_Refv2.0) (Kaur *et al.* 2017) using TopHat v2.1.1 (Trapnell *et al.* 2009) with default parameters and passing the reference annotation with the-G option. Paired-end and single-end reads were mapped separately, and the BAM files were used to calculate the read counts for each gene using featureCounts (Liao *et al.* 2014) (Table S1.1 and Table S1.2). For each sample, the sum of read counts at each gene locus was taken by adding the read counts from the paired-end BAM file and single-end BAM files. Differential expression analysis was carried out on the matrix of read counts for each gene (rows) and sample (columns) using the perl script run_DE_analysis.pl from the Trinity v2.2.0 (Haas *et al.* 2013) suite of programs based on the edgeR method (Robinson *et al.* 2010). Volcano plots were generated for each time-point and differentially expressed genes with false discovery rate (FDR) less than 5% were output for each time-point.

### Functional characterisation and GO enrichment analyses

Gene ontology (GO) enrichment was performed using the Fisher exact test as implemented in R using the topGO package (Alexa & Rahnenfuhrer 2010) with method ‘weight01’ used to adjust for multiple comparisons. Up- and down-regulated DEGs (logFC>2 and FDR<0.05) were considered for the analysis.

### Quantitative reverse-transcriptase PCR (qPCR) analysis

Quantitative reverse transcriptase (qPCR) analysis were performed to study the relative gene expression of *TSub_g7401* (CCT motif protein), *TSub_g3280* (*AGL6*: *AGAMOUS*-like 6), *TSub_g25875* (*AGL20: AGAMOUS*-like 20), *TSub_g20101* (*AGL12: AGAMOUS*-like 12), *TSub_g15483* (*AGL8: AGAMOUS*-like 8), *TSub_g12538* (*FTb2: FLOWERING LOCUS T b2), TSub_g11956* (*WNK5:* With no lysine (K) kinase.) and *TSub_g21607* (*AGL4: AGAMOUS*-like 4) in response to FR-enriched and B-enriched lights. The *Pisum sativum* L. (Alaska) ubiquitin (PUB3) gene L81141.1 was used as reference gene. Leaf tissue was collected at the same time of the day (6 h after lights were switched on) from plants at third-leaf stage grown in both environments (E1 and E2) and used for isolation of total RNA using Spectrum^TM^ Plant Total RNA kit (Sigma). The High-Capacity cDNA reverse transcription kit with RNase inhibitor kit (Appliedbiosystems) was used to synthesise cDNA from 500 ng RNA. Primers for qPCR were designed using NCBI Primer-BLAST (Ye *et al.* 2012) (Table S7). Primer efficiency was determined using LinRegPCR 2014.x software (Ruijter et al. 2009) and was close to the ideal value of 2 for all primer pairs (between 1.8 and 1.9). Reactions were set up with 20 ng cDNA and 3 pmol (each) forward and reverse primers using KiCqStart SYBR Green qPCR Low ROX Ready Mix (Sigma-Aldrich, St Louis, USA), and run on an Applied Biosystems 7500 Fast Real-Time PCR System under the following conditions: 95°C initial denaturation for 30 sec, followed by 40 cycles of 95°C denaturation for 3 sec plus primer annealing/extension at 60°C for 30 sec. Finally, melt-curve analyses were performed (one cycle of 95°C for 15 min, 60°C for 1 min, increasing by 0.1°C min^−1^ to 95°C for 15 sec, 60°C for 15 sec). Data were analysed with the Applied Biosystems 7500 software (version 2.0.6), and values for ΔC_T_ and ΔΔC_T_ calculated according to Schmittgen & Livak (2008). Each sample was run in three technical replicates for each gene.

### Statistical analysis

Differences on gene expression were analysed by one-factor analysis of variance (ANOVA) and the least significant difference (LSD) test at the 5% level of significance.

## RESULTS

### Far-red enriched light accelerates floral onset

To study the effect of light quality on time to flowering, plants were grown simultaneously in controlled environments under the same temperature (24/20 °C day/night) and photoperiod (20 h) regime, but different light spectra. Environment 1 (E1) consisted of a R: FR ratio of 2.89 (FR-enriched light) and Environment 2 (E2) consisted of a R: FR ratio of 5.86 (blue-(B) enriched light). We observed a clear effect of light spectra on floral onset in subterranean clover. Time to flowering was significantly reduced (*P* < 0.001) under FR-enriched light by 14 days compared to B-enriched light (Fig. 1, A and B).

### Gene response to different light spectra

Given the clear effect of the FR-enriched spectrum on the acceleration of floral onset, we aimed to identify the influence of light on gene expression related to flowering induction.

#### Transcriptome dynamics in response to light spectra

RNA-Seq technology was used to analyse variations in gene expression related to changes in the R: FR ratio across three time-points (TP) based on the attainment of precise developmental milestones under E1 growing conditions (low R: FR ratio). This correlated to 14 growing days for TP1 (third-leaf stage), 42 growing days for TP2 (pre-flowering stage under FR-enriched light) and 47 growing days for TP3 (flowering stage under FR-enriched light). From a total of 31,272 protein-coding genes identified in the subterranean clover advanced assembly Tsub_Refv2.0 (Kaur *et al.* 2017), 644 genes were differentially expressed in response to different light spectra in at least one of the three time-points analysed (Table S2). Clear differences in the number of differentially expressed genes (DEGs) were observed in response to the light treatments (Fig. 2B; Fig. 3, A and B; Table S2). The highest number of DEGs was found at TP1 with a total of 448 genes, of which 418 were exclusively differentially expressed at this stage (93%; Table S3). Of the 448 DEGs at TP1, 241 were up-regulated and 207 were down-regulated in B-enriched light compared to FR-enriched light. At TP2, we found 156 DEGs (135 up- and 21 down-regulated) with 122 of them exclusively differentially expressed at this point (78%; Table S4). At TP3, we identified 80 DEGs (16 up- and 64 down-regulated), with 65 of them exclusively differentially expressed at this point (81%; Table S5).

**Figure 3.**
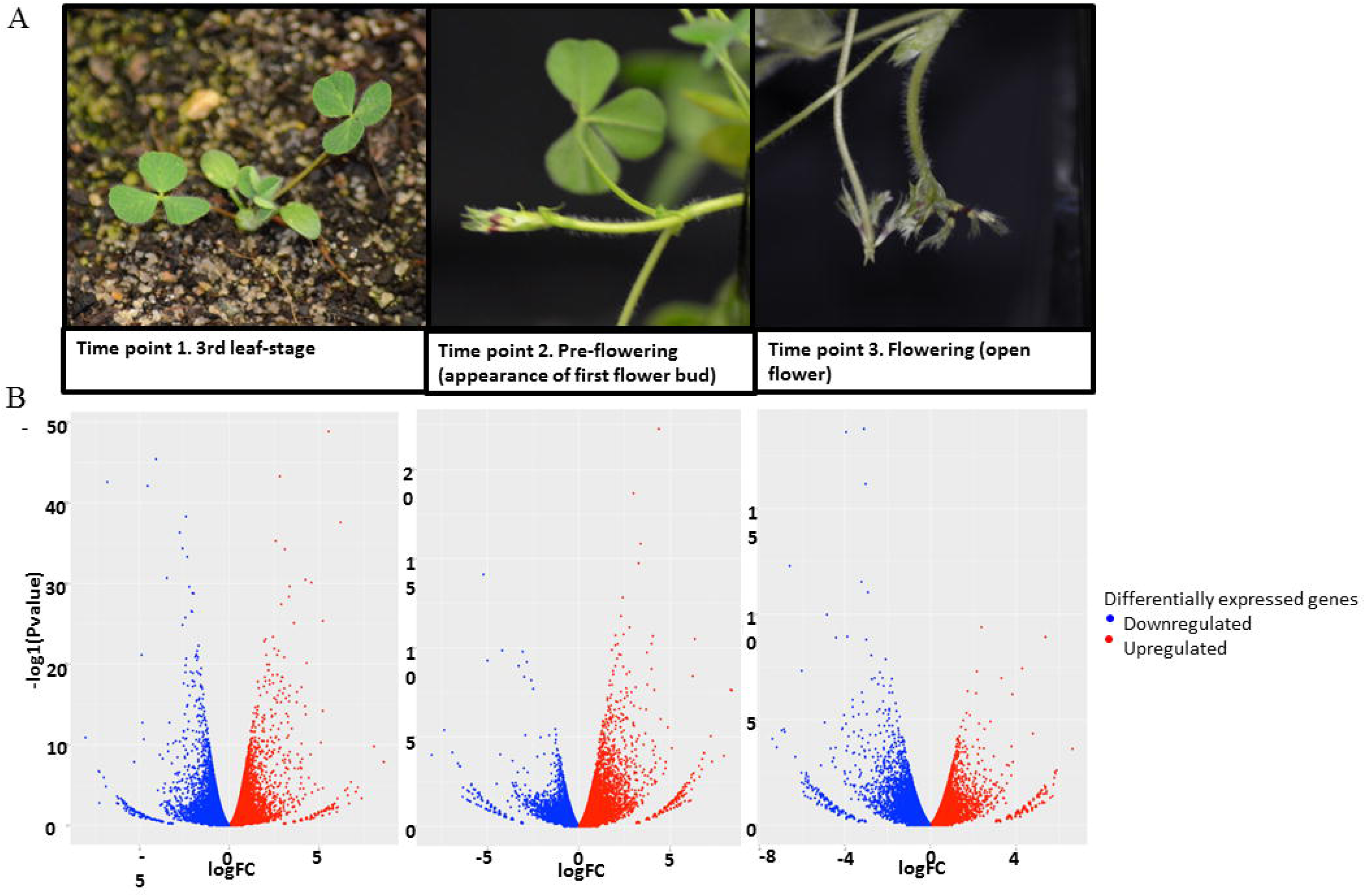
A. The number of genes differentially regulated depending on the light provided. Red represents genes up-regulated and blue represents genes down-regulated under blue-enriched lights at the different time-points (TP) selected in this study when plants in the E1 environment attained the following developmental milestones: third-leaf stage (TP1), appearance of the first flower bud (TP2), open flower (TP3). B. Gene expression differences for genes with FDR < 0.05 and logFC > 2. The Venn diagram presents the number of genes that are exclusively differentially expressed at each developmental stage, with the overlapping regions showing the number of genes that are differentially expressed at two or more developmental stages.

### Far-red and blue-enriched lights affect different biological processes

To identify the major functional biological categories represented by DEGs in response to light spectra, we performed gene ontology (GO) enrichment analysis. Changes in gene expression were seen in the three different time-points in GO terms, suggesting biological processes are differentially affected by light quality. A total of 459 (71.3%) of all DEGs were assigned GO categories corresponding to several aspects of metabolic and cellular processes, response to stress, plant defence, ion localization and transport, and flowering-related processes among others (Table S6a–c).

When analysing DEGs in both environments at TP1, the most significantly enriched GO categories with respect to up-regulated genes (p-value < 0.05) were related to defence response, mainly to biotic stimuli such as bacteria or fungus (Table S6a). Some of those genes were related to flavonoid pathways (Tsub_g874 and Tsub_g13273) and LRR disease resistance related proteins (Tsub_g14165, Tsub_g23725 and Tsub_g6245; Table S3). On the other hand, the GO terms related to ‘ion transport’ and floral development were predominantly associated with down-regulated genes in TP1 (Table S6a). Related to the red light signalling category, a CCT motif protein (TSub_g7401) was down-regulated under B-enriched light. Most of the genes identified in the floral development categories were MAD-box proteins (TSub_g15483, TSub_g25875, TSub_g21607 and TSub_g3280; Table S3).

At TP2, biological processes related to nodulation (GO: 0009877), defence response (GO: 0006952), and vegetative to reproductive transition (GO: 00110228) were up-regulated in B-enriched light compared to FR-enriched light. In contrast, genes involved in the secondary compound biosynthesis of isoflavonoids and flavonoids (GO: 0009717 and GO: 0009813) and anthocyanin-containing compounds (GO: 0009718) were down-regulated (Tables S4 and S6b).

At TP3, genes associated with biological processes related to photoperiodism, flowering (GO: 2000028 and GO: 0048573) and developmental vegetative growth (GO: 0080186) were up-regulated under B-enriched light, and therefore their expression was lower under FR-enriched light (Table S6c). We found that TSub_g17978 was the same gene involved in those biological processes (Table S5). On the other hand, genes associated with the response to stress (GO: 0006950) were down-regulated. Additionally, we found other categories related to stress response: high light intensity (GO: 0009644), response to hydrogen peroxide (GO: 0042542), L-proline biosynthetic process (GO: 0055129), and detection of visible light (GO: 0009584; Table S6c).

### Far-red enriched light promotes the floral induction network

The expression of genes related to flowering promotion across species occurred at an earlier stage of plant development under FR-enriched light than B-enriched light. We identified 13 DEGs related to light signalling, meristem identity and flowering promotion, of which ten were exclusively expressed at TP1, one at TP2 and two at TP3. One of the DEGs was shared between TP1 and TP2 (Table 1). All of the flowering-related DEGs identified at TP1 were down-regulated under B-enriched light, as their expression was higher under FR-enriched light. Of those, we found a gene-coding protein related to light signalling, a CCT motif protein (TSub_g7401), and an uncharacterised protein identified as a probable serine/threonine-protein kinase (with no lysine kinase 5-*WKN5*; TSub_g11956). Additionally, we observed five different MADS-box transcription factors involved in floral promotion (TSub_g15483, TSub_g21607, TSub_g3280, TSub_g25875 and TSub_g20101; Tables I and Table S2). The floral promoter TSub_g12538, which is a possible pea *FLOWERING LOCUS T b2* (*FTb2*), was among the down-regulated genes. Gene-coding proteins potentially linked with floral promotion were also identified: an uncharacterised protein (TSub_g22633), a possible *GIBBERELLIN 20 OXIDASE 2* (*GA20OX2*) involved in the promotion of floral transition, a sugar transporter (TSub_g15042), possible *SUCROSE-PROTON SYMPORTER 2* (*SUC2*), and a homeobox related protein that may be related to meristem identity (TSub_g491; Table 1 and Table S2).

**Table 1.**
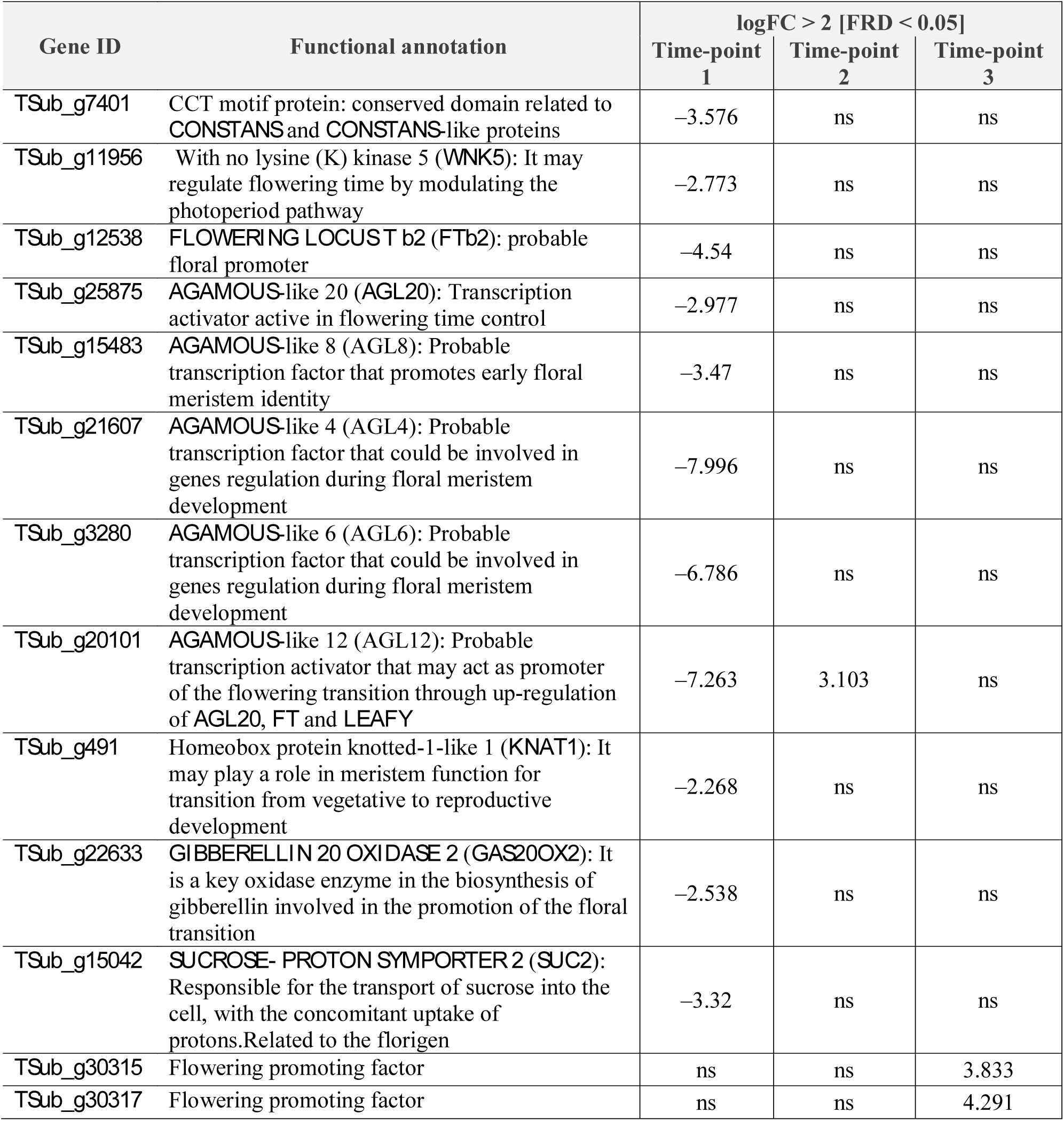
LogFC > 2 [FRD < 0.05] of up- and down-regulated flowering-related genes identified at each time-point when comparing blue-enriched with far-red enriched lights. ns = not significant

| Gene ID | Functional annotation | logFC > 2 [FRD < 0.05] |  |  |
| --- | --- | --- | --- | --- |
|  |  | Time-point 1 | Time-point 2 | Time-point 3 |
| *TSub_g7401* | CCT motif protein: conserved domain related to *CONSTANS* and *CONSTANS*-like proteins | −3.576 | ns | ns |
| *TSub_g11956* | With no lysine (K) kinase 5 (*WNK5*): It may regulate flowering time by modulating the photoperiod pathway | −2.773 | ns | ns |
| *TSub_g12538* | *FLOWERING LOCUS T b2 (Ftb2)*: probable floral promoter | −4.54 | ns | ns |
| *TSub_g25875* | *AGAMOUS*-like 20 (*AGL20*): Transcription activator active in flowering time control | −2.977 | ns | ns |
| *TSub_g15483* | *AGAMOUS*-like 8 (*AGL8*): Probable transcription factor that promotes early floral meristem identity | −3.47 | ns | ns |
| *TSub_g21607* | *AGAMOUS*-like 4 (*AGL4*): Probable transcription factor that could be involved in genes regulation during floral meristem development | −7.996 | ns | ns |
| *TSub_g3280* | *AGAMOUS*-like 6 (*AGL6*): Probable transcription factor that could be involved in genes regulation during floral meristem development | −6.786 | ns | ns |
| *TSub_g20101* | *AGAMOUS*-like 12 (*AGL12*): Probable transcription activator that may act as promoter of the flowering transition through up-regulation of *AGL20*, *FT* and *LEAFY* | −7.263 | 3.103 | ns |
| *TSub_g491* | Homeobox protein knotted-1-like 1 (*KNAT1*): It may play a role in meristem function for transition from vegetative to reproductive development | −2.268 | ns | ns |
| *TSub_g22633* | *GIBBERELLIN 20 OXIDASE 2 (GAS20OX2)*: It is a key oxidase enzyme in the biosynthesis of gibberellin involved in the promotion of the floral transition | −2.538 | ns | ns |
| *TSub_g15042* | *SUCROSE- PROTON SYMPORTER 2 (SUC2)*: Responsible for the transport of sucrose into the cell, with the concomitant uptake of protons. Related to the florigen | −3.32 | ns | ns |
| *TSub_g30315* | Flowering promoting factor | ns | ns | 3.833 |
| *TSub_g30317* | Flowering promoting factor | ns | ns | 4.291 |

All of the DEGs related to flowering induction identified at TP2 and TP3 were up-regulated under B-enriched light. At TP2, one DEG was identified, a MADS-box protein (TSub_g20101). Subsequently, at TP3, we found two DEGs associated with flowering promotion factors (TSub_g30315 and TSub_g30317; Fig. 4 and Table 1).

**Figure 4.**
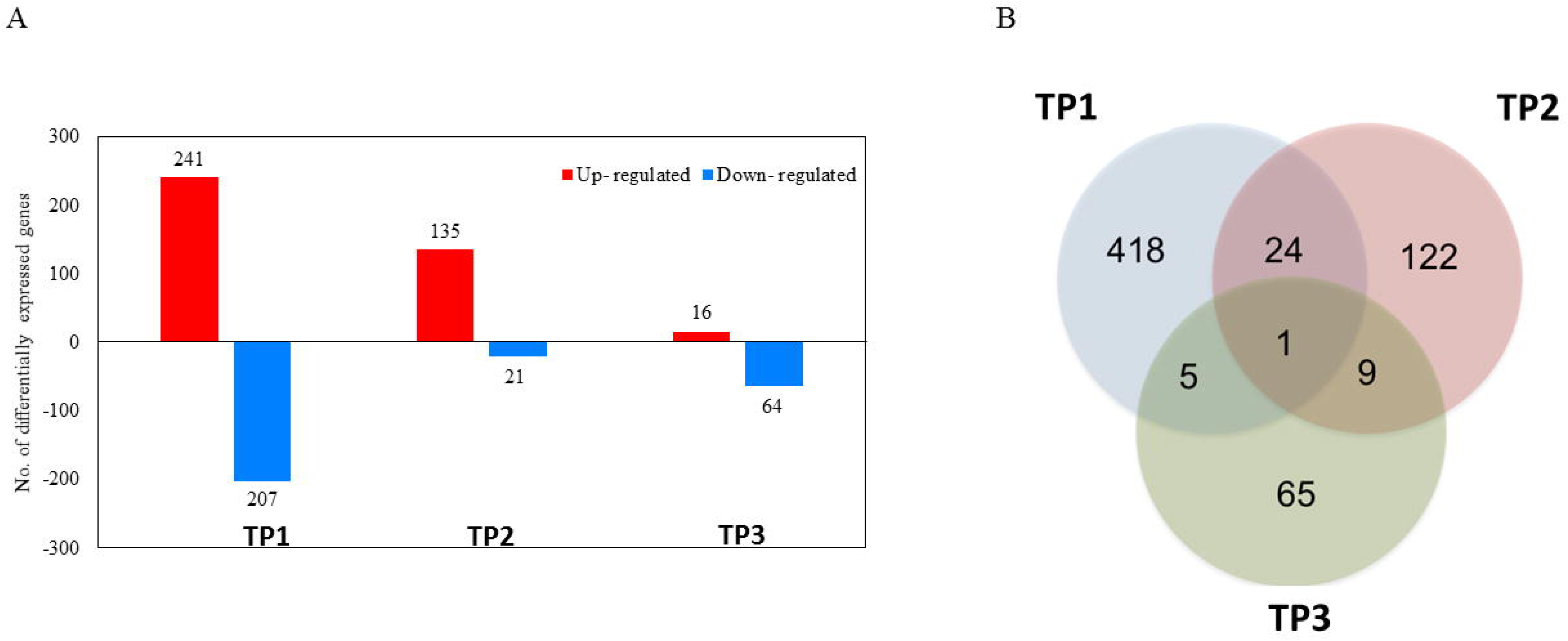
Differentially expressed genes related to flowering induction by different light spectra. A Genes identified at time-point 1 (TP1) when plants reached third-leaf stage, these genes were down-regulated under blue-enriched light (high R: FR). B. Genes identified at time-points 2 (TP2) and 3 (TP3), these genes were up-regulated under blue-enriched light (high R: FR). *CRY:* cryptochrome, *PHYA:* phytochrome *A, PHYB:* phytochrome B, CCT: CCT motif protein, *KNAT1:* homeobox protein knotted-1-like 1, *AGL8: AGAMOUS*-like 8, *FTb2: FLOWERING LOCUS T b2, SUC2:* sucrose-proton symporter 2, *AGL4: AGAMOUS*-like 4, *AGL6*: *AGAMOUS*-like 6, *AGL20: AGAMOUS*-like 20, *AGL12: AGAMOUS*-like 12, *WNK5:* with no lysine (K) kinase, *GAS20OX2*: gibberellin 20 oxidase 2, FP1 and FP2: flowering promoters.

### Expression of flowering genes in response to far-red and blue-enriched lights

The relative expression of flowering induction related genes at the third-leaf stage (TP1) decreased between 1.6 – 104.28 fold changes under B compared to FR-enriched lights. (Fig. 5A). *TSub_g21607*, a MADS-box transcription factor involved in floral promotion, showed the highest fold change (−104.28) compared to the other genes studied. The rest of MADS-box presented lower fold changes, between −1.60 and −20.79, together with *TSub_g7401* (−14.16 change-fold) and *TSub_g12538* (−4.09).

**Figure 5.**
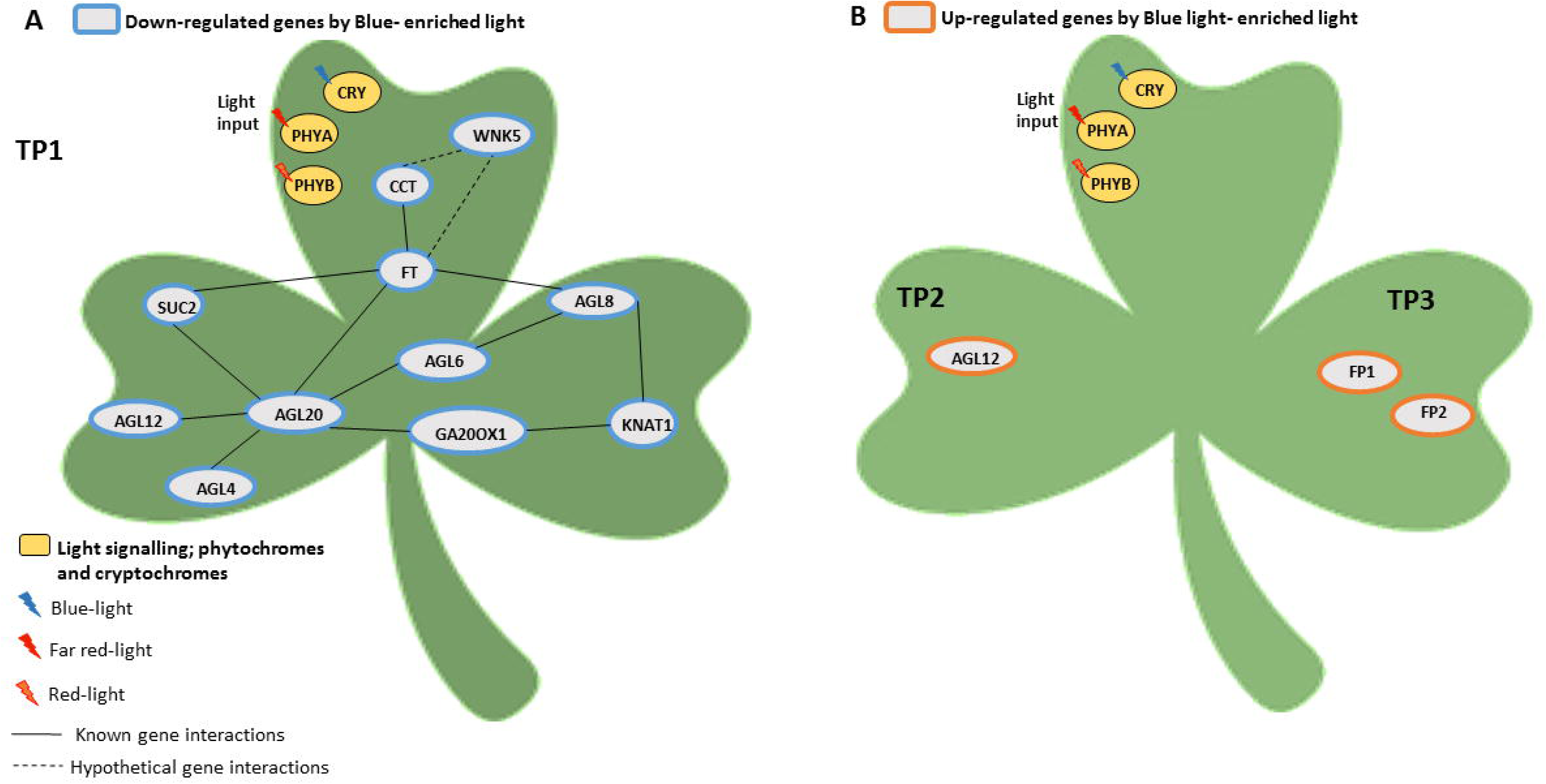
Relative gene expression of *TSub_g7401* (CCT motif protein), *TSub_g3280* (*AGL6*: *AGAMOUS*-like 6), *TSub_g25875* (*AGL20: AGAMOUS*-like 20), *TSub_g20101* (*AGL12: AGAMOUS*-like 12), *TSub_g15483* (*AGL8: AGAMOUS*-like 8), *TSub_g12538* (*FTb2: FLOWERING LOCUS T b2), TSub_g11956* (*WNK5:* With no lysine (K) kinase) and *TSub_g21607* (*AGL4: AGAMOUS*-like 4) in response to Far Red-enriched (FR) and Blue-enriched (B) lights. Gene expression was quantified by qPCR at third-leaf stage (14 d after sowing). Data is represented as expression of the gene of interest relative to *Pisum sativum* L. (Alaska) ubiquitin (PUB3) gene L81141.1 as reference gene using the 2^−ΔCt^ method (Schmittgen & Livak 2008). Values are means ± SE (n = 3 replicates of 2 plants).

Relative expression (2^−ΔCt^) of the studied genes was significantly higher in response to FR-enriched light (Fig.5B). Two genes showed the biggest response to light quality, *TSub_g21607*, a MADS-box transcription factor involved in floral promotion and *TSub_g11956*, a WNK related to the photoperiod pathway.

## DISCUSSION

For the first time, we have demonstrated growing plants under FR light accelerates the up-regulation of genes related to floral initiation pathways using the long-day plant subterranean clover as a model. We identified novel genes which link photoperiod and FR-light signalling in addition to the up-regulation of genes related to plant defence under B-enriched light.

RNA-Seq time-course analysis identified 13 DEGs related to light signalling, meristem identity and flowering promotion. There was clear evidence in the transcriptome of up-regulation of gene networks related to flowering at a very early stage of plant development (TP1, third-leaf stage) when plants were grown under FR-enriched light. Under B-enriched light, the up-regulation of this network was not expressed until TP2 and TP3 when the expression of *AGL12* and floral promoting factors was identified. These results are in accordance with the acceleration of time to flowering observed under FR-enriched light (low R: FR ratio; 14 days faster) compared to B-enriched light (high R: FR ratio).

The genetic link for the integration of photoperiod, light perception and circadian clock pathways has been well characterised in Arabidopsis and involves the B-box transcription factor *CO*. The floral promoter *CO* protein is up-regulated by long-day growing conditions and its expression is stabilised by FR light through *PHYTOCHROME A* activity (Kim *et al.* 2008; Pin & Nilsson 2012; Song *et al.* 2013). In Medicago, a temperate legume as subterranean clover, it has been suggested that the integration of responses to day length and light quality may not be regulated by *CO*-like (*COL*) genes (Wong *et al.* 2014; Weller & Ortega 2015). In contrast, in subterranean clover, we identified the expression of a CCT motif (*TSub_g7401*) at an early developmental stage (TP1) under FR-enriched light. That result was supported by qPCR analysis. We found that the relative expression (2^−ΔCt^) of *TSub_g7401* gene, homologous to a conserved domain related to *CO* and *COL* proteins, was higher under the FR compare to the B-enriched light at the same 20h photoperiod with a −14.16 fold change. Therefore our results suggest that, in subterranean clover, at same photoperiod conditions (20 h) FR enriched light induces *COL* gene expression. In addition, we identified a serine/threonine-protein kinase (TSub_g11956), *WNK5* known to be involved in regulating time to flowering in response to photoperiod in Arabidopsis. Wang *et al.* (2008) reported that *WNK5* acts to delay flowering in Arabidopsis. In our case, the higher relative expression of *TSub_g11956* was observed under FR-enriched light, it is suggested that this homologous gene in subterranean clover is related to flowering induction and its expression is modulated by light quality. Our results suggest floral initiation under FR-enriched light is mediated by CO-like and WNK-like proteins. To our knowledge, a direct interaction between WNK-like protein and *FTb2* has not previously been reported. For subterranean clover under FR-enriched light, the relative expression of *TSub_g11956* (homologous of *WNK*-like gene) was higher than for *TSub_g7401* (CCT motif related to *CO* and *COL*), so although a direct interaction can be inferred, further work is suggested to elucidate changes in pattern expression throughout the day.

The CO protein has a role in activating the expression of FT (Kim et al. 2008). FT-like proteins from several species function similarly to FT with respect to the induction of flowering, transport in phloem, and interaction with FLOWERING LOCUS D-like proteins (Hecht et al. 2011). In our experiment, at the first time-point (TP1) under FR-enriched light, we found a gene-coding protein (TSub_g12538) homologous of a pea FTb2 whose relative expression was also higher under FR compared to B-enriched lights. This protein is expressed specifically in pea and Medicago leaves under long days and meets the characteristics of the classical ‘florigen’ (Weller & Ortega 2015). The differential expression of this homologous gene may suggest its role as a floral promoter in subterranean clover.

Downstream in the photoperiod pathway, we identified four MADS-box transcription factors exclusively expressed at TP1 under FR-enriched light (TSub_g15483, TSub_g21607, TSub_g3280, and TSub_g20101). We also identified one MAD-box TF (*AGL*12) that was up-regulated at TP1 under FR-enriched light and not identified until TP2 under B-enriched light (TSub_g25875). These TFs have been identified as key components of the networks that control the transition from vegetative to reproductive stages and flower development in Arabidopsis (Tapia-Lpez *et al.* 2008). These flowering promoters are homologous to *AGAMOUS-LIKE* (*AGL*) MADS-box proteins: *AGL4, AGL6, AGL8, AGL12* and *AGL20*. *AGL8* promotes floral determination in response to FR-enriched light (Hempel *et al.* 1997). *AGL20* has been described as an integrator of the vernalisation, autonomous and photoperiod pathways controlling flowering in Arabidopsis (Lee *et al.* 2000). It has also been associated with gibberellins in the induction of flowering (Moon *et al.* 2003). *AGL12* has been described as an important floral promoter through the up-regulation of A*GL20* (Tapia-López *et al.* 2008). The expression of *AGL4* and *AGL6* is associated with gene regulation during floral meristem and floral organ development, with both found mainly in flowers (Ma *et al.* 1991; Pelaz *et al.* 2000; Dreni & Zhang 2016). The fact that *AGL12* was found at TP1 under FR-enriched light and at TP2 under B-enriched light provides further evidence that FR-enriched spectra accelerate flowering induction.

In addition to the DEGs related to the photoperiod flowering pathway and downstream cascade, we identified a further three DEGs related to flowering at TP1 under FR-enriched light: TSub_g491, TSub_g22633 and TSub_g15042. The TSub_g491 gene-coding protein is a homolog of the *KNAT1* protein and involved in the development of both vegetative and reproductive meristems (Scofield *et al.* 2007; Aguilar-Martinez *et al.* 2015). The TSub_g22633 is homologous to *GAS20OX2*, which is involved in the promotion of floral transition in Arabidopsis (Andrés *et al.* 2014). The TSub_g15042 is a sugar transporter, *SUC2* involved in the transport of *FT* through phloem companion cells in leaves to the meristem for the induction of floral organ formation (Corbesier *et al.* 2007). At the later developmental time-points (TP2 and TP3), only three genes were exclusively differentially expressed under B-enriched light. At TP2, we identified a gibberellin-related protein (TSub_15579) homologous to *AT5G14920*, which is shown to be related to flowering induction in Arabidopsis over-expressing the *ZEITLUPE/LOV KELCH PROTEIN 1*, a blue light photoreceptor (Saitoh *et al.* 2015). At TP3, we found two genes (TSub_g30315 and TSub_g30317) described as homologs of flowering-promoting factor-like proteins in Medicago. The fact that genes related to reproductive processes were identified at TP1 under FR-enriched light and at TP2 and TP3 under B-enriched light is in accordance with the accelerated flowering observed under a low R: FR ratio.

Interestingly, we found contrasting responses to light quality related to reproduction and defence mechanisms. In our experiment, at TP1 and TP2, growing plants under B-enriched light (high R: FR ratio) enhanced the expression of genes involved in flavonoid and anthocyanin pathways. At TP3, we found down-regulation of genes related to the L-proline biosynthetic process, which is involved in the response to abiotic stress (Devlin 2016). This is in agreement with studies indicating plant health responses are modulated by B-enriched and FR-enriched light. For example, B-enriched light can enhance the production of secondary metabolites such as flavonoids and anthocyanins, which are defensive compounds against fungus, bacteria and environmental stresses (Johkan *et al.* 2010). Similarly, plants grown under FR-enriched light (low R: FR ratio) can express a weak defence phenotype (Cerrudo *et al.* 2012; Ballaré 2014). Our findings provide further support that the stress response is active under FR-enriched light, which may be due to the preferential allocation of resources to reproduction over defence.

The results from this study will contribute to a deeper understanding of the molecular basis of flowering and the possible negative correlation between reproductive and defence mechanisms in response to different light quality in long-day plants. Using RNA-Seq and tightly controlled light spectra, we have identified key genes which link photoperiod and FR-light signalling coding for a CCT motif, a WNK-like and an *FTb2*-like protein, and the active downstream cascade. The earlier down-regulation and the lower relative expression of these genes observed under B-enriched light compared to FR-enriched light may explain their role in the acceleration of floral onset.

## ACKNOWLEDGMENTS

This work was supported by the grant-in-aid funding awarded to Dr Kaur by the AW Howard Memorial Trust Inc. We thank Mr R. Creasy, Mr B. Piasini and Mr L. Hodgson for glasshouse expertise and Ms S. Wells for technical assistance. We acknowledge the supercomputing resources provided by the Pawsey Supercomputing Centre with funding from the Australian Government and the Government of Western Australia.

## ACCESSION CODE

The RNA-Seq sequencing data has been made available (under embargoed) at https://zenodo.org (DOI: 10.5281/zenodo.1184974).

